# Chromosome-level genome assemblies for two quinoa inbred lines from northern and southern highlands of Altiplano where quinoa originated

**DOI:** 10.1101/2024.06.10.598385

**Authors:** Yasufumi Kobayashi, Hideki Hirakawa, Kenta Shirasawa, Kazusa Nishimura, Kenichiro Fujii, Rolando Oros, Giovanna R. Almanza, Yukari Nagatoshi, Yasuo Yasui, Yasunari Fujita

## Abstract

Quinoa, an annual allotetraploid plant native to the Andean highlands of South America, is emerging as an important seed crop for global food and nutrition security due to its ability to grow in marginal environments and its excellent nutritional properties. Because quinoa is partially allogamous, we have developed quinoa inbred lines necessary for molecular genetic analysis. Our comprehensive genomic analysis showed that the quinoa inbred lines fall into three genetic subpopulations: northern highland, southern highland, and lowland. Lowland and highland quinoa are the same species, but have very different genotypes and phenotypes. Lowland quinoa has relatively small grains and a darker grain color, and is widely tested and grown around the world. In contrast, the white, large-grained highland quinoa is grown in the Andean highlands, including the region where quinoa originated, and is exported worldwide as high-quality quinoa. Recently, we have shown that viral vectors can be used to regulate endogenous genes in quinoa, paving the way for functional genomics of quinoa. However, although a high-quality assembly has recently been reported for a lowland quinoa line, genomic resources of the quality required for functional genomics are not available for highland quinoa lines. Here we present high-quality chromosome-level genome assemblies for two highland inbred quinoa lines, J075 representing the northern highland line and J100 representing the southern highland line, using PacBio HiFi sequencing and dpMIG-seq. The assembled genome sizes of J075 and J100 are 1.29 and 1.32 Gb, with contigs N50 of 66.3 and 12.6 Mb, and scaffold N50 of 71.2 and 70.6 Mb, respectively, comprising 18 pseudochromosomes. The repetitive sequences of J075 and J100 represent 72.6% and 71.5% of the genome, the majority of which are long terminal repeats (*Gypsy* and *Copia*), representing 44.0% and 42.7% of the genome, respectively. The *de novo* assembled genomes of J075 and J100 were predicted to contain 64,945 and 65,303 protein-coding genes, respectively. The high quality genomes of these highland quinoa lines will facilitate quinoa functional genomics research on quinoa and contribute to the identification of key genes involved in environmental adaptation and quinoa domestication.

## 1 Introduction

The composition of global staple crops has remained largely unchanged for decades. However, with the rapid pace of climate change, crop diversification will be critical to securing the future of our food supply (Mayes et al., 2012; Massawe et al., 2016; Krug et al., 2023). To date, approximately 2,500 plant species in 160 families have been domesticated, but less than 300 of these species are available on the global market. The majority of the world’s population relies on only a handful of plant species for their caloric intake (Dirzo and Raven, 2003; Meyer and Purugganan, 2013; Massawe et al., 2016). Among crops considered underutilized and relatively neglected, quinoa (*Chenopodium quinoa* Willd.) is in the spotlight due to its excellent nutritional properties and remarkable ability to grow under harsh environmental conditions (Masawe et al. 2016)

Quinoa is a C3 annual plant belonging to the Amaranthaceae family, which also includes crops such as sugar beet (*Beta vulgaris* L.) and spinach (*Spinacia oleracea* L.). Both its seeds and leaves are consumed as food. Notably, quinoa seeds boast an exceptional nutritional profile, offering a well-rounded balance of the five essential macronutrients: proteins, fats, carbohydrates, vitamins, and minerals (Navruz-Varli and Sanlier, 2016; Nowak et al., 2016; Vilcacundo and Hernández-Ledesma, 2017; Rodriguez et al., 2020). Specifically, quinoa is rich in high-quality proteins, encompassing all nine essential amino acids, such as lysine, which the human body cannot produce on its own (Ruales and Nair, 1992; Burrieza et al., 2019; Dakhili et al., 2019). Additionally, it is gluten-free, making it a safe option for individuals with gluten allergies (Peñas et al., 2014). Quinoa leaves, too, are a valuable source of essential amino acids, vitamins, and minerals (Pathan et al., 2019). Furthermore, quinoa contains various functional components, such as bioactive peptides, polysaccharides, saponins, polyphenols, flavonoids, and phytoestrogens, which are believed to possess antioxidant and anti-inflammatory properties, as well as potential benefits for hypertension and diabetes management (Hirose et al., 2010; Vilcacundo and Hernández-Ledesma, 2017; Ren et al., 2023). In quinoa, the dispersal unit (i.e., seed) is botanically classified as an achene—a type of fruit containing a single seed encased in a dry, non-opening outer shell, referred to as the pericarp (Prego et al., 1998; Burrieza et al., 2019). The seed features a curved embryo positioned around the edges, enveloping a central area known as the perisperm or basal body, which provides a unique structural characteristic to quinoa seeds (Prego et al., 1998; Burrieza et al., 2019). Unlike other grains like wheat and rice, where essential nutrients are mainly located in the endosperm and outer hull, the valuable amino acids and functional compounds in quinoa are found in the kernel (Hemalatha et al., 2016; Motta et al., 2017; Motta et al., 2019).

Quinoa has been cultivated in the Andes region of South America for over 7,000 years (Dillehay et al., 2007). Revered as the ‘mother grain’ in the pre-Columbian era, it was a fundamental part of the Andean indigenous peoples’ diet, along with llamas and tubers (González et al., 2015; Miller et al., 2021). However, following the Spanish conquest, quinoa’s cultivation and use in indigenous ceremonies were banned, contrasting with the global spread of other Andean crops like tomatoes (*Solanum lycopersicum* L.) and potatoes (*Solanum tuberosum* L.) (Gomez-Pando, 2015). Since the 1950s, quinoa has served as an indicator plant for identifying plant virus species due to its susceptibility to various plant viruses that cause local lesions (Uschdraweit, 1955; Hein, 1957; Yasui et al., 2016). Recognizing its potential, the National Academy of Sciences (NAS) and the National Research Council (NRC) highlighted quinoa as one of the underexploited crops with significant economic promise (NAS, 1975; NRC, 1989). The National Aeronautics and Space Administration (NASA) has also explored quinoa as a nutritious food source for astronauts on extended space missions (Schlick and Bubenheim, 1993; 1996). The UN’s designation of 2013 as the “International Year of Quinoa” aimed to underscore its potential in enhancing food and nutrition security and supporting sustainable agriculture (Bazile et al., 2016). Recently, although Bolivia and Peru remain the leading producers, accounting for nearly 80% of the world’s supply, quinoa is being researched and grown in over 120 countries (Alandia et al., 2020).

Quinoa can be grown at temperatures from below freezing to near 40°C, at low to high latitudes, and from lowlands along the coast to highlands above 4,000 m, and is highly tolerant to abiotic stresses such as drought, high salt, low temperatures, hail, and frost (Jacobsen, 2003; Hariadi et al., 2011; Zurita-Silva et al., 2014; Yasui et al., 2016; Mizuno et al., 2020). For example, the area around Salar de Uyuni, the world’s largest salt flat, located in Bolivian Altiplano (3,800 m above sea level), where annual precipitation is less than 200 mm and saline-alkaline soil prevails, has been known as a production area for high-quality quinoa, which can only be grown as a crop in this harsh environment (Bonifacio, 2019). However, cultivating quinoa outside the Andean region presents challenges such as its low tolerance to high temperatures, increased vulnerability to pests and diseases, and sensitivity to day-length variations that need to be addressed (Gomez-Pando et al., 2019).

Quinoa is an allotetraploid species with a chromosome count of 2*n* = 4*x* = 36 and an estimated genome size of approximately 1.5 Gb, comprising two sub-genomes, A and B (Palomino et al., 2008; Yangquanwei et al., 2013). For years, the complex genome of quinoa, a consequence of its allotetraploidy, along with genetic diversity stemming from partial outcrossing due to the presence of both hermaphrodite and female flowers on the same plant, had posed challenges to molecular analysis (Maughan et al., 2004; Christensen et al., 2007). To overcome this situation, our collaborative group (Yasui et al., 2016) and two subsequent independent groups (Jarvis et al., 2017; Zou et al., 2017) decoded the quinoa genome with the technology available at the time. Following this, we have developed more than 130 genotyped quinoa inbred lines tailored for molecular studies, elucidating genotype-phenotype relationships for salt stress responses and important growth traits (Mizuno et al., 2020). Our comprehensive genomic analyses, employing single-nucleotide polymorphisms (SNPs), revealed that these quinoa inbred lines can be categorized into three genetic sub-populations: the northern highland, southern highland, and lowland groups (Mizuno et al., 2020). Furthermore, we have established a method to analyze the function of endogenous genes in quinoa by using virus-induced gene silencing (VIGS) and virus-mediated overexpression (VOX), paving the way for functional genomics analysis (Ogata et al., 2021).

Accurate genome sequencing is crucial for advancing functional genomics in quinoa, shedding light on its unique properties, and tackling the challenges it faces in both local and global production. Thus far, genome assemblies have been produced for four quinoa lines: two from lowland lines, Kd (Yasui et al., 2016) and QQ74 (Jarvis et al., 2017), and two from southern highland lines, Real (Zou et al., 2017) and CHEN125 (Bodrug-Schepers et al., 2021). Although these assemblies are fragmented and do not fully reflect the chromosomal biology of quinoa, the chromosome-level assembly QQ74 V2 has recently been reported for the lowland quinoa line QQ74 (Rey et al., 2023). However, unlike lowland quinoa lines, no useful genomic information is available for highland quinoa lines: there are still no reports of genome assemblies of northern highland quinoa lines or chromosome-level assemblies of southern highland quinoa lines. In particular, lowland and highland quinoa lines are the same species, but differ significantly not only in genotype but also in phenotype, for example, highland quinoa lines are difficult to grow in lowland areas, while lowland quinoa lines are difficult to grow in highland areas (Mizuno et al., 2020). Northern highland quinoa lines have traditionally been grown around Lake Titicaca in Peru and Bolivia, where quinoa is believed to originate. The quinoa lines that have been attempted to be grown outside the Andean region such as in Europe, the United States, Asia, and Africa, are basically lowland quinoa lines with relatively small grains and a darker color, while the white, large-grained quinoa lines grown in the Andean region of South America and exported to outside the Andean region such as in Europe, the United States, and Asia are highland quinoa lines. The Altiplano highlands of Peru and Bolivia, situated at elevations of 3,000 to 4,000 m, are among the world’s largest quinoa-producing regions, supplying high-quality, organically cultivated quinoa globally. However, due to the harsh conditions of this environment, they are particularly vulnerable to climate change (Bonifacio, 2019). Given these circumstances, a reference genome assembly that provides accurate genomic information on highland quinoa are essential to develop effective breeding strategies to address these challenges and to understand the process of quinoa domestication, including its adaptation to harsh environments and its origins.

In this study, we present chromosome-level genome assemblies for two inbred lines of highland quinoa: J075, a representative of the northern highland, and J100, a representative of the southern highland (Mizuno et al., 2020; Ogata et al., 2021). These assemblies were achieved by integrating PacBio HiFi long-read sequencing with the dpMIG-seq method, which is a multiplexed inter simple sequence repeat genotyping by sequencing, using degenerate oligonucleotide primers (Nishimura et al., 2024). The reference genomes obtained in this study will provide the basis for advancing functional genomics in quinoa to facilitate the development of climate-adapted highland quinoa breeding materials and contribute to a better understanding of the domestication process of quinoa, including its adaptation to harsh environments and its origin. These findings have the potential to provide clues for improving various crops to make them more adaptable to climate change.

## 2 Materials and methods

### 2.1 Plant materials and growth conditions

Quinoa inbred lines, derived through single-seed descent via self-crossing, were cultivated in phytotrons as described previously (Yasui et al., 2016; Mizuno et al., 2020; Ogata et al., 2021). In this study, we used J075 as a representative of the northern highland lines, J100 as a representative of the southern highland lines, and J082 as a representative of the lowland quinoa lines (Mizuno et al., 2020). The J075 and J100 plants were grown until they produced more than a dozen fully expanded leaves, under conditions of 25 ± 5°C and a 12-hour light/12-hour dark photoperiod. To assess the color phenotype of the J075, J100, and J082, they were grown for eight weeks under the same conditions as stated above.

### 2.2 Processing data for the whole genome sequence of quinoa

High molecular weight DNA was extracted from fully expanded leaves of the J075 and J100 lines, using a single plant from each line and the Genomic-tip Kit (Qiagen). SMRT sequencing libraries were prepared with the SMRTbell Express Template Prep Kit 2.0 (PacBio, Menlo Park, CA, USA) and sequenced on a PacBio Sequel II system, utilizing two SMRT Cell 8M units. The PacBio subreads were converted to HiFi reads using the circular consensus sequencing (CCS) program (http://github.com/PacificBiosciences/ccs). HiFi reads from each quinoa line were assembled using Hifiasm (Cheng et al., 2021). Chromosome-scale pseudomolecules were then created using the reference-guided scaffolding software, RagTag ver. 2.1.0 (Alonge et al., 2022).

An F_6_ mapping population (n =183) was generated from a cross between the quinoa inbred lines J100 and J027. Genomic DNA was extracted from the young leaves of each line. Using genomic DNA from the two parental lines and 183 F_6_ plants, the dpMIG-seq library was constructed as described previously (Nishimura et al., 2024). The libraries were sequenced using an Illumina HiSeq X sequencer. Low quality and adapter sequences were removed using Trimmomatic ver. 0.38 (Bolger et al., 2014) with the settings ‘HEADCROP:17 SLIDINGWINDOW:4:20 LEADING:20 TRAILING:20 MINLEN:25’. The resultant high quality reads were mapped onto hifiasm-assembled J100 contigs using BWA ver. 0.7.17 (Li, 2013) and sorted using samtools ver. 1.18 (Danecek et al., 2021), and then generated a vcf file by mpileup in bcftools ver. 1.16 (Danecek et al., 2021). A total of 13,526 SNPs were uniformly selected for genetic linkage map construction in software Lep-MAP3 ver. 0.5 (Rastas, 2017). In detail, the markers were identified as informative through the ParentCall2 module and anchored to LGs comparable to the contigs in the pseudomolecule assembled by RagTag using the SeparateChromsomes2 module by adjusting the LOD values (ranging from 15 to 40). Next, a total of 29,751 SNPs were extracted as anchor markers from the 151 contigs comprising these scaffolds and linkage maps were reconstructed for each contigs in each scaffold using LepMap3 as described above. The high-density genetic linkage maps were visualized using ALLMAPS pipeline (Tang et al., 2015).

The genomic structures of J075 and J100 were compared to QQ74 (v2, id60716) from CoGe (https://genomevolution.org/coge/) using D-GENIES (Cabanettes and Klopp, 2018), with alignments performed by Minimap2 ver. 2.26 (Li, 2021). Using the same methods, the genomic structures of Real (Zou et al., 2017) and CHEN125 (Bodrug-Schepers et al., 2021) were compared to J100 genome assembly in this study.

### 2.3 Assessing assembly completeness

The completeness of the genome assemblies for J075 and J100 was assessed using the Benchmarking Universal Single-Copy Orthologs (BUSCO) database, ver. 5.3.2 (Simao et al., 2015), in genome mode. This assessment searched for genes conserved in embryophyte species. The data sets, embryophyta_odb10, created on 10 September 2020, include 50 genomes and 1614 BUSCOs, respectively. The completeness of the assembled genomes for J075 and J100 was further evaluated using the Long Terminal Region (LTR) Assembly Index (LAI) (Ou et al., 2018), which assesses genome quality based on intergenic genome information using LTR retrotransposons (LTR-RT). LTR retrotransposon candidates were identified using LTRharvest and LTR_FINDER_parallel and were further classified using LTR_retriever ver. 2.9.6 (Ou and Jiang, 2018). The continuity of repeat space in the assemblies was evaluated by the LAI, calculated using the LTR_retriever package.

### 2.4 Repetitive sequence detection and genome functional annotation

*De novo* repeat libraries for the pseudomolecule sequences J075 and J100 were constructed using RepeatModeler ver. 2.0.3 (Flynn et al., 2020). Repetitive sequences were identified and classified using these de novo repeat libraries along with RepeatMasker ver. 4.1.2 (Flynn et al., 2020). Putative protein-coding genes in the genome sequences were predicted using the BRAKER2 pipeline, which incorporates protein evidence via GeneMark-ES (Bruna et al., 2021). Proteins were aligned to the genome using ProtHint (Bruna et al., 2020), which integrates the Spaln (Iwata and Gotoh, 2012; Gotoh et al., 2014) and DIAMOND (Buchfink et al., 2015) protein aligners, generating specific gene model parameters. These parameters were then applied in AUGUSTUS ver. 3.4.0 (Hoff and Stanke, 2019) for gene prediction. Within the BRAKER pipeline, Viridiplantae protein sequences from the OrthoDB (Kriventseva et al., 2019) served as the evidence protein dataset.

Homology-based gene annotation was conducted using GeMoMa ver. 1.9 (Keilwagen et al., 2019), utilizing genome, gene annotation, and mapped RNA-seq data from quinoa reference lines (Yasui et al., 2016; Rey et al., 2023). This homology-based gene prediction by GeMoMa was complemented by comparison with *ab initio* gene annotation by BRAKER, with GeMoMa modules being used to obtain the final gene annotation. The quality of putative protein-coding genes was evaluated using the EnTAP functional annotation package ver. 0.10.8 (Hart et al., 2020), with curated databases from UniProtKB/Swiss-Prot (https://ftp.uniprot.org/pub/databases/uniprot/current_release/knowledgebase/complete/uniprot_sprot.fasta.gz), UniProtKB/TrEMBL (https://ftp.uniprot.org/pub/databases/uniprot/current_release/knowledgebase/complete/uniprot_trembl.fasta.gz), and NCBI’s Plant Protein (https://ftp.ncbi.nlm.nih.gov/refseq/release/plant/) for similarity searches (DIAMOND E-value < 0.00001). These searches were followed by alignment to the eggNOG database ver. 4.5 using eggNOG-mapper ver. 2.1 (Huerta-Cepas et al., 2016; Cantalapiedra et al., 2021). Gene family associations provided the basis for Gene Ontology (GO) term assignments, identification of protein domains (Pfam) (Finn et al., 2014), and associated pathways (KEGG) (Kanehisa et al., 2014). Genome circular plots were created using Circa (https://omgenomics.com/circa/).

### 2.5 Orthology prediction and phylogenomic analysis

Orthologous cluster, phylogenetic, and gene family evolution analyses of quinoa J075 and J100, along with six other flowering species (*Arabidopsis thaliana*, *Oryza sativa*, *Zea mays*, *Amaranthus hypochondriacus*, *Beta vulgaris*, and *C. quinoa* QQ74), were conducted using the OrthoVenn3 platform (Sun et al., 2023). The OrthoFinder algorithm identified orthologous gene families across these eight plants, which included quinoa lines partitioned by two subgenomes. Clustered gene families were visualized using UpSetR (Conway et al., 2017), which facilitated the analysis of overlapping families among multiple plants and highlighted unique clusters within each species. Overlapped and unique gene families were used for GO enrichment analysis in the platform.

The phylogenetic and gene family contraction and expansion analysis were automatically run on the platform. A total of 2,137 single-copy orthologs from these plants were aligned using MUSCLE, with conserved sequences subsequently trimmed using TrimAl. Phylogenetic trees were then constructed using the maximum likelihood method, based on the JTT+CAT evolutionary model implemented in FastTree2. In addition to these analyses, CAFE5 was employed to detect gene family expansion and contraction by extrapolating species gene family sizes and evolutionary timelines. Divergence times among lineages containing each quinoa genome were estimated using the TimeTree5 database (http://www.timetree.org/), with divergence events for *A. thaliana* vs *O. sativa*, *A. hypochondriacus* vs *O. sativa*, *B. vulgaris* vs *O. sativa*, *A. thaliana* vs *A. hypochondriacus*, *A. thaliana* vs *B. vulgaris*, and *A. hypochondriacus* vs *B. vulgaris* occurring at 160, 160, 160, 118, 118, and 49 mya, respectively. GO enrichment analysis was also automatically performed on the platform to explore the functional attributes of the expanded and contracted gene families as mentioned above.

### 2.6 Genome-wide replication analysis

The synonymous substitution rate (*K*s) distributions between the two sub-genomes of quinoa, *C. pallidicaule* (A-progenitor genome, v4.0, id53872) (Mangelson et al., 2019) and *C. suecicum* (B-progenitor genomes, v2, id52047) (Rey et al., 2023) from CoGe (https://genomevolution.org/coge/) were calculated by wgd ver. 2.0.26 (Chen and Zwaenepoel, 2023). For the calculation of *K*s distributions for one-to-one orthologs, the dmd module was utilized to extract orthologs via an all-versus-all BLASTP search using the DIAMOND algorithm. Subsequently, the ksd module was employed to construct the one-to-one ortholog *K*s distributions among the genomes.

### 2.7 Genome synteny analysis

Syntenic gene pairs among quinoa lines were identified using the MCScan pipeline implemented in python [https://github.com/tanghaibao/jcvi/wiki/MCscan-(Python-version)] (Tang et al., 2008). Within the pipeline, the “jcvi.compara.catalog ortholog” function was used to search for syntenic regions. The “jcvi.compara.synteny screen” function was employed to identify macro-syntenic blocks, which are defined as having more than 30 collinear genes within a syntenic block. The “jcvi.compara.synteny mcscan” function was utilized for micro-synteny analysis at the gene level. Finally, the “jcvi.graphics.karyotype” function was used to visually display each synteny as a karyotype.

For identification of structural variants (SVs) calling using SyRI (ver.1.6) (Goel et al., 2019), minimap2 (ver. 2.24) (Li, 2021) was used to generate pairwise genome alignments with parameters ‘-ax asm5’. The alignment result was subsequently passed to SyRI, and SVs consisting of insertions, deletions, inversions, and translocations were kept using default parameters.

## 3. Results

### 3.1 Chromosome-level genome assemblies for southern and northern highland inbred quinoa lines

Based on the results of previous population structure and phenotypic analyses (Mizuno et al., 2020; Ogata et al., 2021), we selected J075 as a representative line of northern highland quinoa and J100 as a representative line of southern highland quinoa. The whole genome sequences of northern highland J075 and southern highland J100 were obtained using PacBio Sequel II sequencing platform, yielding 59.9 Gb and 38.9 Gb of data, respectively (Supplementary Table 1). The PacBio data obtained for J075 and J100 were assembled into 155 and 963 primary contigs, respectively (Supplementary Table 2). For J075, the HiFi-based contig sequences totaled 1.29 Gbp with an N50 of 66.3 Mbp, and for J100, they totaled 1.32 Gbp with a contig N50 of 12.6 Mbp (Supplementary Table 2). Next, we performed reference-guided scaffolding of the HiFi-based contigs, using RagTag in conjunction with the quinoa reference genome (QQ74, v2, id60716) (Rey et al., 2023), which includes all 18 chromosomes of quinoa (Chromosomes 1A to 9A and 1B to 9B). Subsequently, 30 contigs of J075 and 151 contigs of J100 were each mapped to the 18 chromosomes of quinoa (Cq1A-Cq9A and Cq1B-Cq9B; Supplementary Table 3). RagTag analysis resulted in total scaffold lengths of 1.28 Gbp for J075 and 1.27 Gbp for J100, with average scaffold lengths of 71.0 Mbp and 70.7 Mbp, respectively (Supplementary Table 3).

The 30 contigs of J075 could be mapped directly to the reference genome sequence, but the mapping of the 151 contigs of J100 was unclear in some places, so scaffolding was performed using a linkage map created from the J100 x J027 (Mizuno et al., 2020) mapping population. Initially, SNPs were extracted from all contigs assembled using HiFi reads to serve as anchor markers for linkage analysis. In the linkage groups, we confirmed that the majority of contigs mapped to the scaffold assembled by RagTag. Consequently, 29,751 SNPs were extracted as anchor markers from the 151 contigs comprising these scaffolds for linkage analysis. The contigs were then mapped to a linkage group, covering 3,205 cM, using these SNPs (Figure 1 and Supplementary Table 4). Dot plot analysis conducted with D-GENIES showed a significant improvement in assembly continuity for the J100 assembly compared to the previously published genomic scaffolds for the Real line (Zou et al., 2017) and the CHEN125 line (Bodrug-Schepers et al., 2021) (Supplementary Figure 1). These findings indicate that the J100 assembly represents the most contiguous, highest-quality assembly of southern highland quinoa lines to date.

**FIGURE 1.**
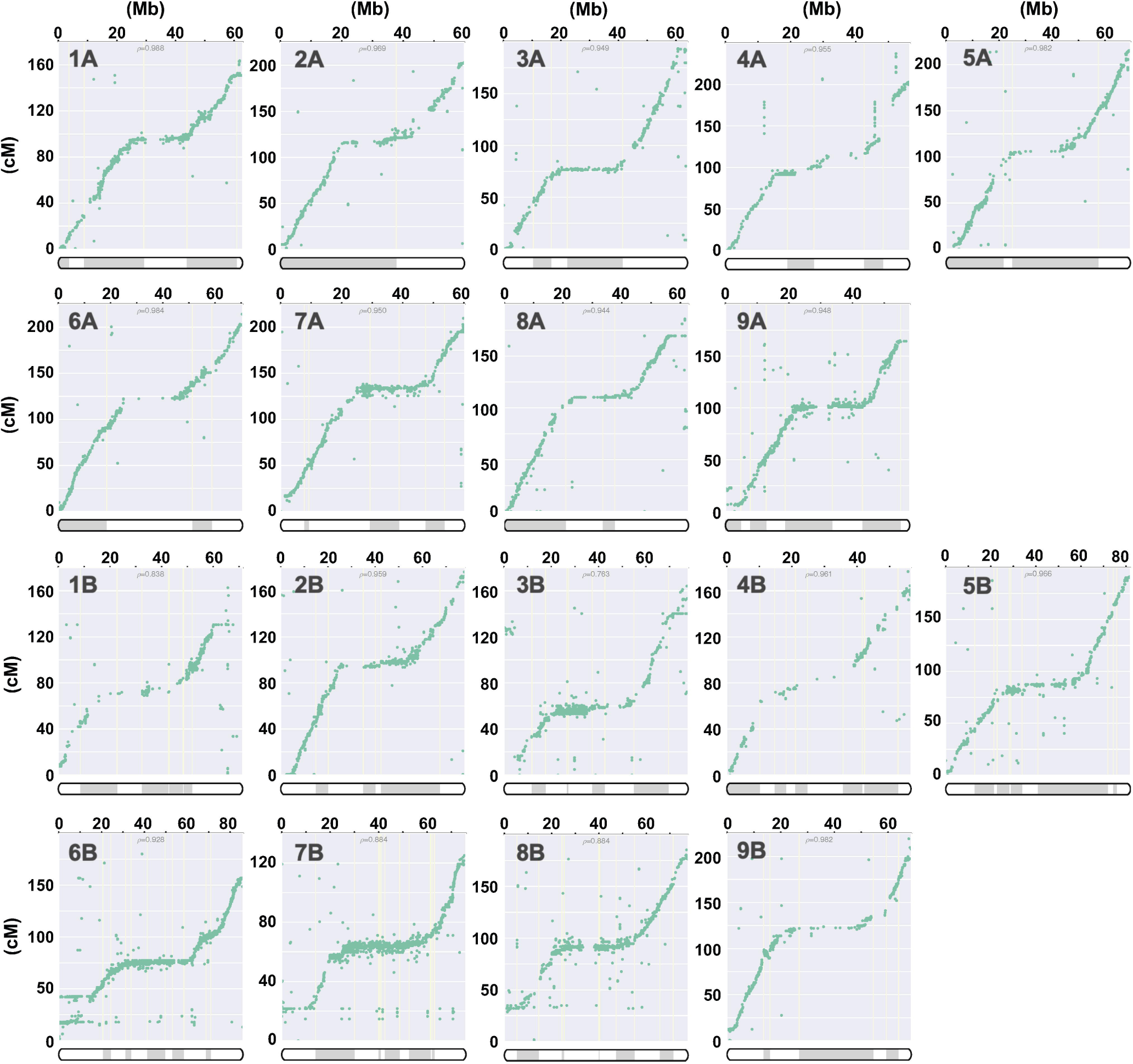
Correlation analysis for quinoa line J100 between linkage groups and the scaffolds, illustrating the physical positions on the pseudochromosomes (x-axis) versus the map locations (y-axis). The pseudochromosomes of the quinoa J100 homology-based assembly scaffold, spanning Chromosomes 1A to 9A and 1B to 9B, have been reconstructed from the genetic map. The p-value represents the Pearson correlation coefficient.

Comparing the pseudomolecule lengths of both J075 and J100 lines and reference genome line QQ74, the N50 lengths of J075 and J100 were both 1.06 times longer than those of QQ74 (Table 1). Dot plot analysis using D-GENIES, comparing the assemblies of J075 and J100 with the reference genome line QQ74, revealed structural variations in Cq7A and Cq3B (Figure 2 and Supplementary Figure 2). For the order of physical positions of contigs in the scaffolds obtained with Hifiasm and RagTag, contigs that were clearly inconsistent with their positions on the linkage map were reassembled according to the order of the linkage map (Supplementary Table 4). These results show that the J075 and J100 assemblies exhibit superior genome contiguity compared to the QQ74 assembly of the reference genome.

**FIGURE 2.**
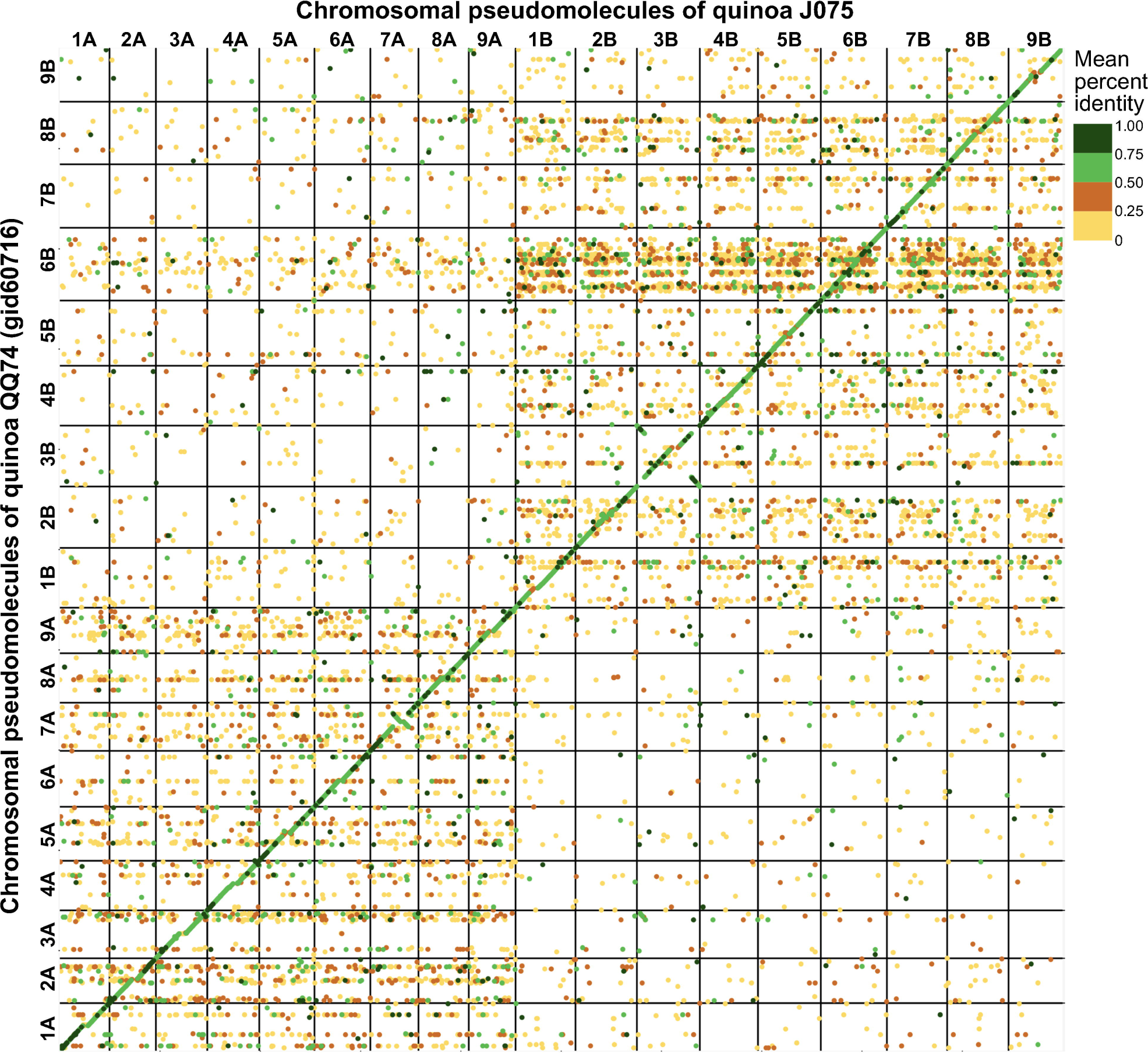
Dotplot image comparing the quinoa J075 pseudomolecules with the reference chromosomes. Dotplot analysis was conducted using minimap2 aligner, and the result was subsequently visualized on the D-GENIES platform (https://dgenies.toulouse.inra.fr). The colors in the dotplot represent identity levels.

**Table 1.** Scaffolding statistics of quinoa J075 and J100 after assembly using RagTag or in combination with Lep-MAP3.

| Statistics | J075 | J100 |
| --- | --- | --- |
| N50(bp) | 71,226,514 | 70,609,007 |
| L50 | 9 | 9 |
| N90(bp) | 59,638,252 | 59,948,908 |
| L90 | 17 | 17 |
| Number of scaffolds | 143 | 813 |
| Gaps | 12 | 149 |
| N count (bp) | 1,200 | 14,900 |
| Largest scaffold (bp) | 84,258,211 | 86,085,297 |
| Smallest scaffold (bp) | 28,991 | 18,494 |
| Average scaffold length (bp) | 9,041,347 | 1,623,460 |
| Total length (bp) | 1,292,912,647 | 1,319,872,862 |
| Percentage in 18 largest scaffolds | 98.9 | 96.4 |

### 3.2 Genome annotation and gene prediction

The completeness of the genome assembly for all scaffolds and non-localized contigs was quantified using the BUSCO database. We used the embryophyta_odb10 dataset, which contains 1,614 core genes. Scaffolds and non-localized contigs of J075 and J100 contained a higher number of benchmarking genes (99.2% and 99.1%, respectively), and almost all of those (89.7% and 89.3%, respectively) were duplicated as isoform genes, according to our BUSCO analyses (Supplementary Table 5). The benchmarking gene preservation of J075 and J100 was almost equivalent to that of reference genome line QQ74. The completeness of these genome assemblies was further validated using the LAI that evaluates assembly continuity using LTR-RTs. A higher LAI score indicates better quality of repetitive and intergenic sequence spaces due to the identification of more intact LTR-RTs. J075 and J100 presented relatively higher LAI scores, at 17.40 and 17.75, respectively (Supplementary Table 6). Based on these quality assessment results, the J075 and J100 genomes were classified as being of reference quality based on the assembly of repetitive and intergenic sequence spaces (draft quality, LAI < 10; reference quality, LAI between 10 and 20; gold quality, LAI > 20). We then analyzed repetitive and transposable sequences in J075 and J100. The repetitive sequences comprised 938.9 Mb (72.6%) and 944.0 Mb (71.5%) of the pseudomolecule sequences of J075 and J100, respectively (Supplementary Table 6). The most common LTR-RTs superfamily, *Gypsy* and *Copia*, comprised 44.0% and 42.7% of the genomes of J075 and J100, respectively (Figure 3). These results indicated that both J075 and J100 contain a similar proportion of repetitive and transposable sequences. The GC contents ratio of the two genomes averaged 37% across the genome length, but this percentage was lower in the central region of most chromosomes, where a region containing an unknown LTR retrotransposon was detected (Figure 3 and Supplementary Figure 3).

**FIGURE 3.**
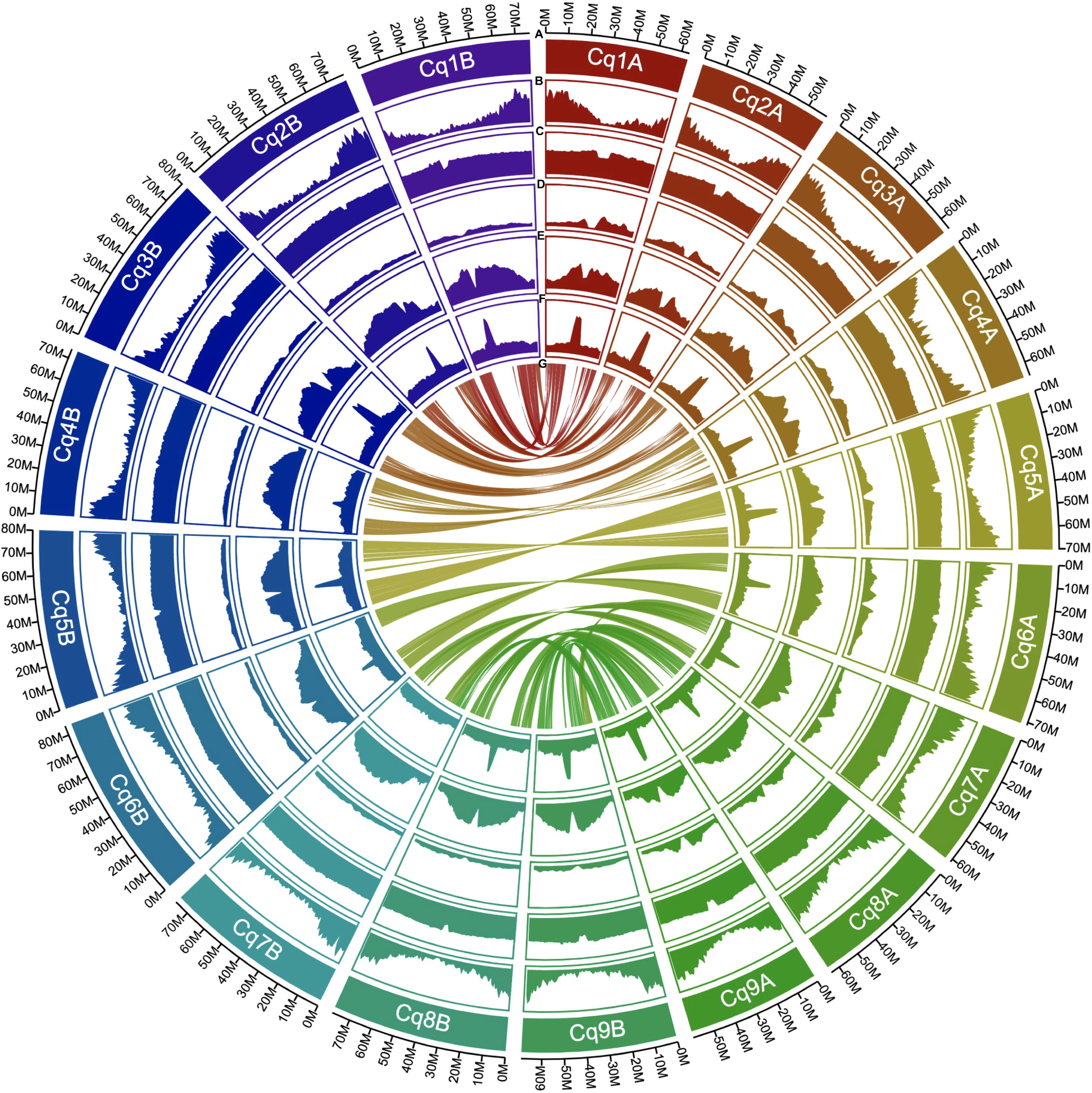
Distribution of genomic features in quinoa J075. From the outermost to the innermost, the tracks represent the following: assembled 18 pseudochromosomes **(A)**, gene density **(B)**, GC content **(C)**, copia-type retrotransposon fragments **(D)**, gypsy-type retrotransposon fragments **(E)**, unknown retrotransposon fragments **(F)**, and intergenomic collinearity **(G)**.

Next, gene models for the J075 and J100 genomes were developed using a comprehensive approach that included *ab initio* and homology-based predictions. Initially, BRAKER2 and GeMoMa predicted a total of 100,169 and 97,309 genes, respectively, from the softmasked scaffolds of J075 and J100 using protein evidence. These genes underwent further analysis, including more sensitive DIAMOND searches against the UniProtKB and NCBI NR databases and domain searches against the Pfam database. Genes containing transposon-related keywords were excluded from the similarity search results. Furthermore, genes filtered by the ‘Viridiplantae’ and ‘Eukaryotes’ categories in the EggNOG Tax Scope were classified as ‘best’ genes. As a result, 65,303 and 64,945 genes from J075 and J100, respectively, were classified as ‘best’ genes (Supplementary Table 7). These predicted genes were found to be densely distributed at the terminal ends of chromosomes (Figure 3).

### 3.3 Gene family analysis and genome evolution in quinoa

To explore the evolutionary history of the quinoa gene family, orthologous gene families were identified using the protein sequences of J075 and J100 predicted in this study, along with those from six other flowering species (*A. thaliana*, *O. sativa*, *Z. mays*, *A. hypochondriacus*, *B. vulgaris*, and *C. quinoa* QQ74). Given that quinoa is allotetraploid, their protein sequences were differentiated into the A and B genomes. In total, 27,158 and 27,791 genes from the A and B genomes of J075 (representing 99.4% and 99.5% of the total, respectively) clustered into 26,989 and 27,663 gene families. Similarly, 27,014 and 27,347 genes from the A and B genomes of J100 (accounting for 99.5% of the total each) clustered into 26,873 and 27,197 gene families. Among these, 8,986 gene families were shared across all genomes from the eight plants (Figure 4). We also discovered that the A genomes of J075 and J100 have 28 and 27 unique gene families (with 104 and 101 unique genes, respectively), respectively, while the B genomes have 23 and 24 unique gene families (with 124 and 108 unique genes, respectively) (Supplementary Figure 4 and Supplementary Table 8). Classification of these unique orthologous gene families revealed that nucleic acid metabolic processes and defense response-related GO terms are enriched across multiple sub-genomes in quinoa (Supplementary Table 9).

**FIGURE 4.**
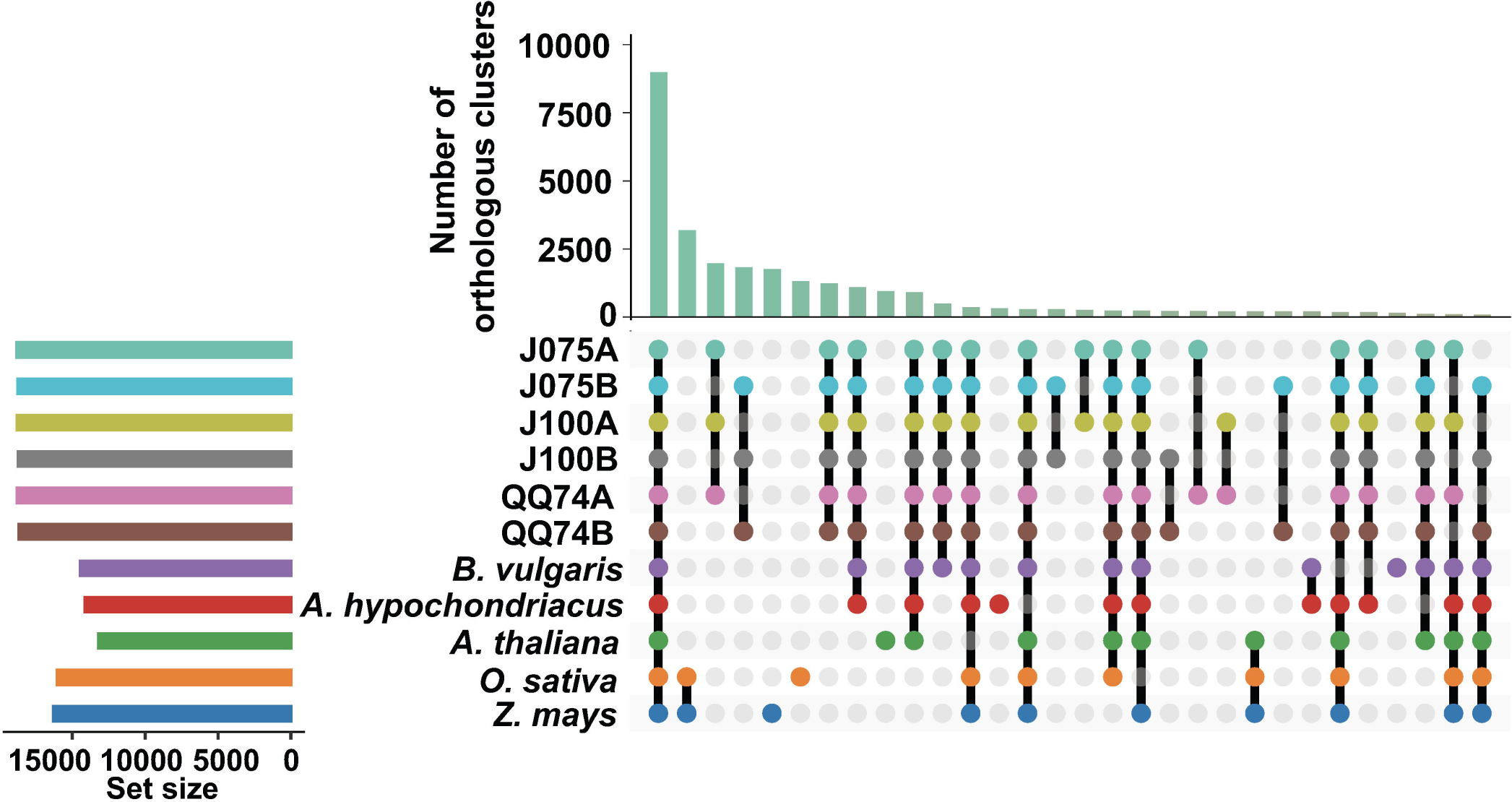
Orthologous cluster analysis of several plant species, including the A and B genomes of quinoa. An UpSet table displays both unique and shared orthologous clusters among the species. The horizontal bar chart on the left shows the number of orthologous clusters per species, while the vertical bar chart on the right indicates the number of clusters shared among the species. Lines represent intersecting sets. The results of the intersecting combinations of J075, J100, QQ74, *B. vulgaris* and *A. hypochondriacus* from this dataset are illustrated as Venn diagrams in Supplementary Figure 4.

We constructed a high-confidence phylogenetic tree and estimated the divergence times of these plant species based on the amino acid sequences from 2,137 single-copy gene families (Figure 5). As expected, the phylogenetic trees generated using concatenated methods confirmed that the A and B genomes of quinoa were divided into two separate clades. Subsequently, the expansion and contraction of orthologous gene families were analyzed using CAFE5. In the lineage leading to the highland quinoa population, 85 and 62 gene families were expanded, while 91 and 63 gene families were contracted in the A and B genomes, respectively. In J075 and J100, 94 and 81 gene families were extended, respectively, which is fewer than the 142 and 161 extended gene families found in the reference quinoa’s A and B genomes (Figure 5 and Supplementary Tables 10 and 11). To investigate the effects of gene gain and loss on quinoa population development, we conducted GO enrichment analysis for the significantly changed gene families in each quinoa line (Supplementary Tables 10 and 11). GO terms common to two or more lines included nucleic acid and protein metabolic processes, as well as defense response-related terms. These results mirrored those specific to quinoa lines in a comparative analysis of orthologous gene families among plant species. Regarding the unique extended gene families, the top five enriched GO terms in J075A were rRNA base methylation, gibberellic acid-mediated signaling pathway, regulation of intracellular pH, tRNA methylation, and multicellular organismal development; in J075B, they were (-)-secologanin biosynthetic process, ATP synthesis coupled electron transport, protein-chromophore linkage, aerobic electron transport chain, and lipid transport; in J100A, they were stress response to copper ion, cell surface receptor signaling pathway, DNA replication, microsporogenesis, and unidimensional cell growth; and in J100B, they were cellular aromatic compound metabolic process, positive regulation of hydrogen peroxide metabolic process, maintenance of chromatin silencing, anatomical structure morphogenesis, and auxin-activated signaling pathway. These expanded and contracted gene families are suggested to reflect trait differences in the J075 and J100 lines.

**FIGURE 5.**
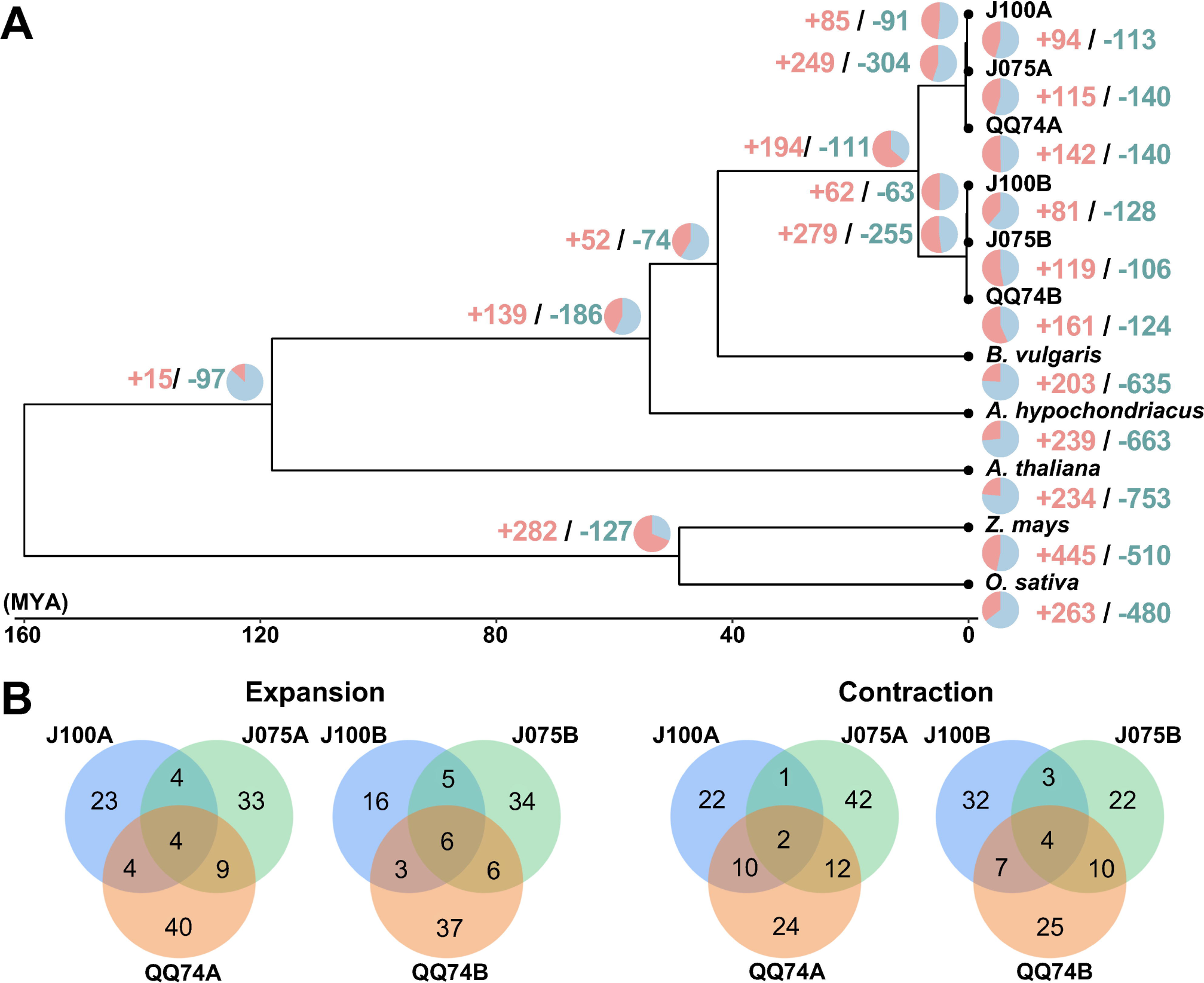
Gene family expansion and contraction in quinoa. **(A)** Phylogenetic tree displaying divergence times and the evolution of gene family size across seven species. The numbers of expanded and contracted gene families are indicated by red and blue numbers, respectively. **(B)** Venn diagrams illustrating the expanded and contracted gene families among J075, J100, and the reference line.

*K*s distributions in the allotetraploid quinoa genome displayed a peak not observed in the ancestral diploid species, suggesting that whole genome duplication may have been caused by hybridization of the ancestral species (Jarvis et al., 2017). Therefore, we calculated the *K*s distribution for one-to-one orthologous genes between both genomes of quinoa and their progenitor genomes. By comparing the divergence between the A- and B-progenitor genomes (J075A and J075B, J100A and J100B, and A- and B-progenitors), we found that the *K*s values in the highland quinoa lines J075 and J100 were consistent with those between the A and B progenitor genomes, as well as with those reported for the previously studied lowland quinoa line QQ74 (Figure 6). The results suggest that hybridization of the A and B genomes occurred simultaneously in the highland lines as it did in the lowland line.

**FIGURE 6.**
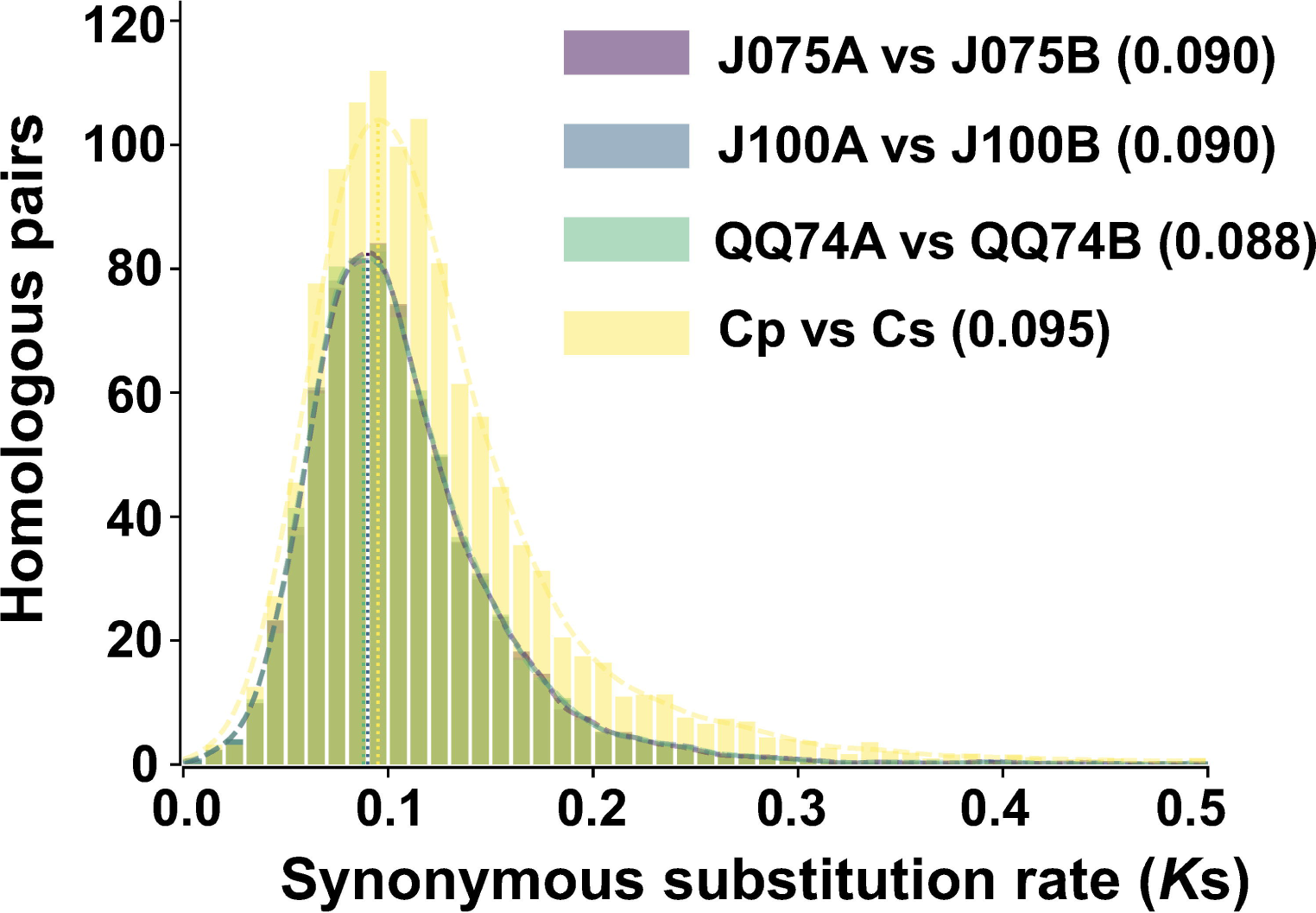
Distribution of synonymous substitution rate for one-to-one orthologous gene sets between A and B genomes of quinoa lines. The comparison between *C. pallidicaule* (Cp) and *C. suecicum* (Cs), as progenitors of the A and B genomes, is also included. Peak values of the density plots for each comparison are indicated in parentheses. In the plots comparing quinoa lines, the colors representing each line overlap.

### 3.4 Synteny analysis and structural-variant detection in quinoa genomes

The collinearity of chromosomes was assessed by comparing the sequences of J075, J100, and reference lines. Based on JCVI-filtered information, we provided the distribution of collinear genes across the chromosomes. The number of collinear blocks in the J075-QQ74, J100-QQ74, and J075-J100 pairs were 71, 70, and 51, respectively, encompassing 44,966, 45,143, and 53,429 genes. These results demonstrated a clear one-to-one syntenic relationship, with overall gene synteny being largely conserved (Figure 7). Despite the high level of synteny across all 18 chromosomes, SyRI identified structural rearrangements when comparing the J075 genome to the J100 genome. Specifically, we identified 100 inversions ranging from 217 bp on chromosome 9B to 9.2 Mb on chromosome 5A (Figure 8 and Supplementary Table 12).

**FIGURE 7.**
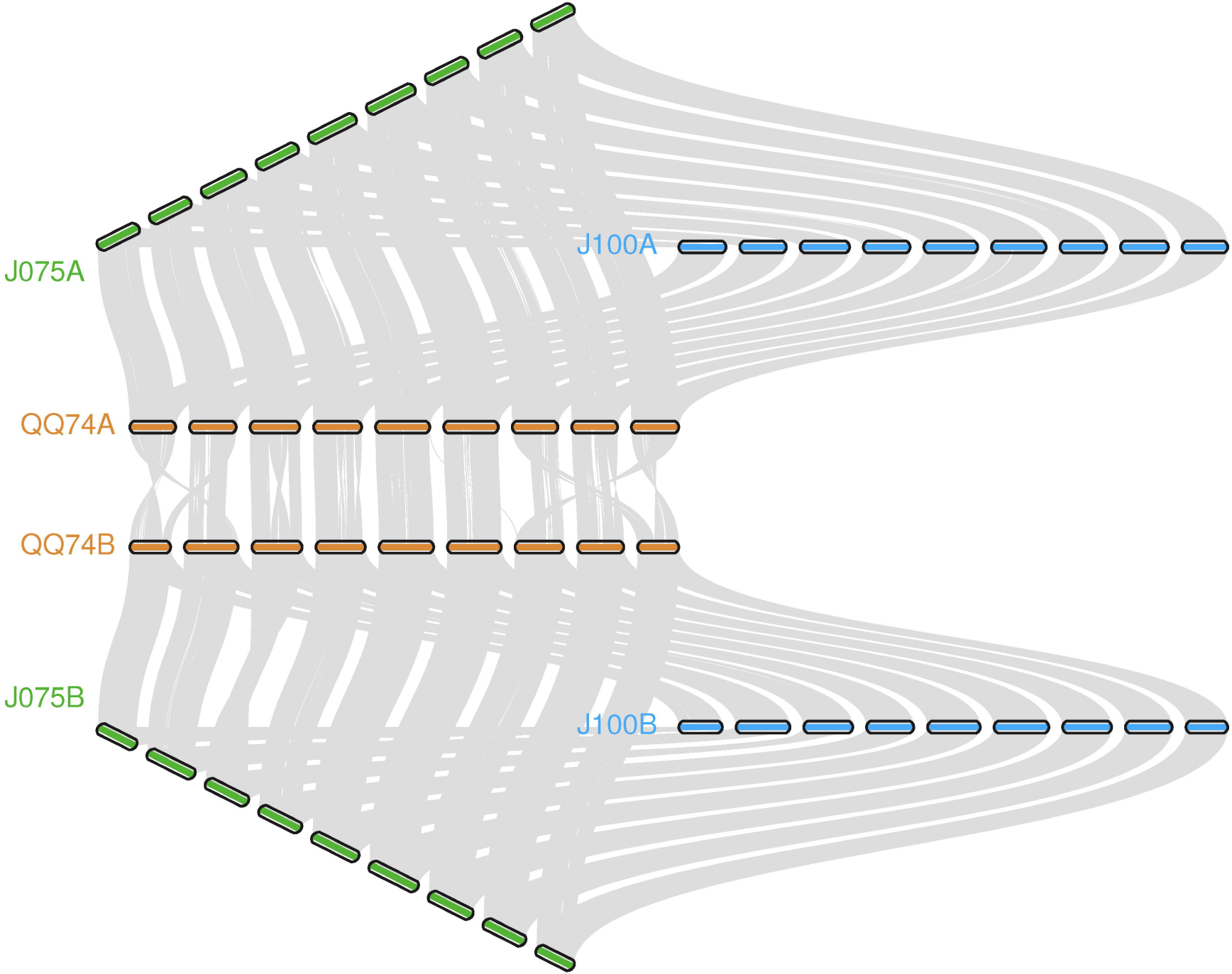
Illustration of the synteny between J075, J100, and the reference line, showcasing conserved genes linked by lines between A and B subgenomes. Syntenic blocks are connected by grey lines.

**FIGURE 8.**
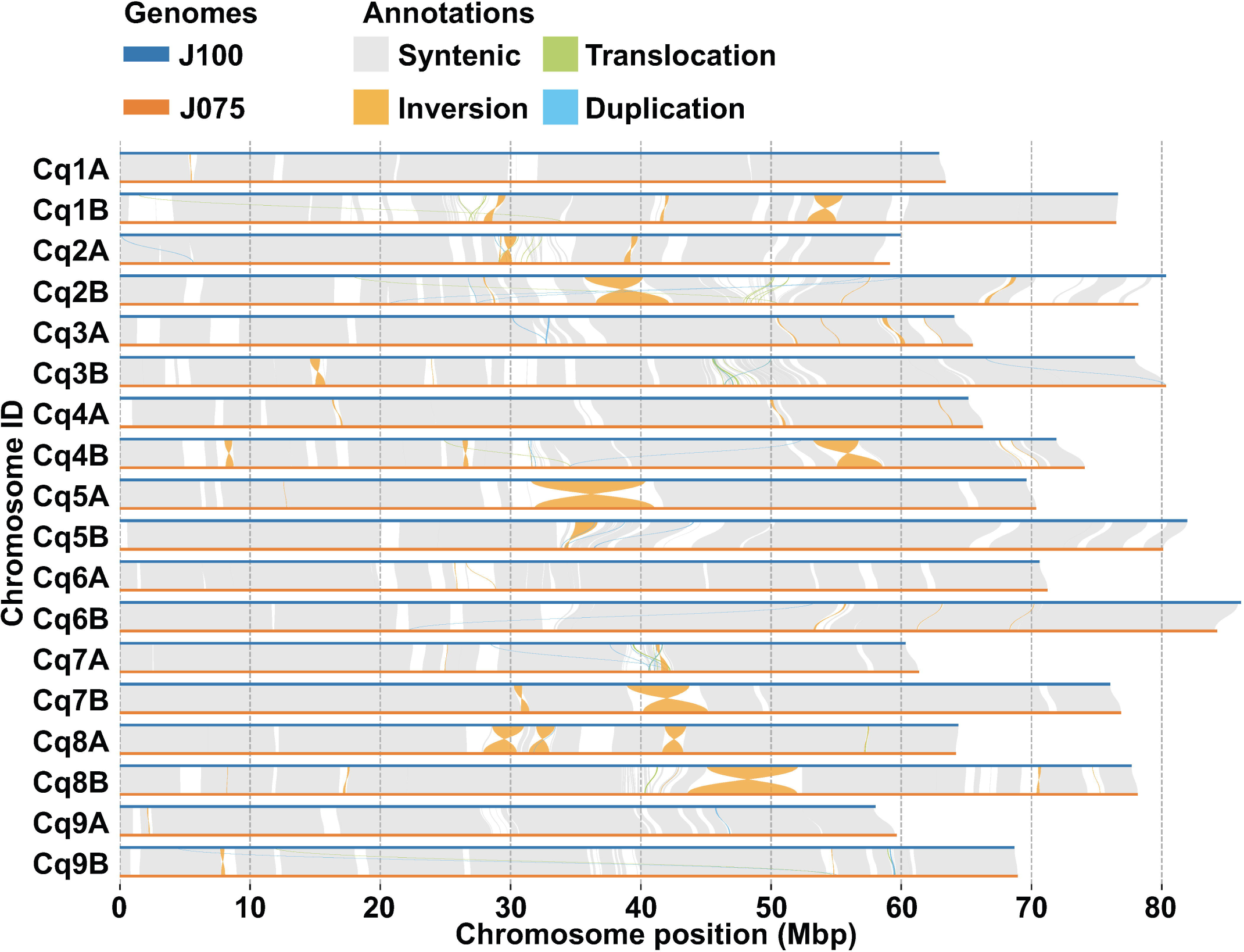
Structural variant detection between J075 and J100. This figure identifies synteny (shown in gray) and structural rearrangements, including inversions, translocations, and duplications, between the chromosomes of J100 (top) and J075 (bottom).

### 3.5 Variation of betalain biosynthesis gene family in highland quinoa lines

We have been working on phenotypic and genotypic associations, such as responses to salt stress and key growth traits, among genetically classified subpopulations of the quinoa inbred lines produced by our research group (Mizuno et al., 2020). Although synteny is retained in most genomic regions among quinoa lines, localized regions with disrupted synteny may harbor genetic factors responsible for their phenotypic differences. A biosynthetic pathway in quinoa that uses L-tyrosine as a precursor to synthesize betanin or amaranthin has been previously reported (Imamura et al., 2019; Ogata et al., 2021) (Supplementary Table 13). We selected genes homologous to the reported betalain biosynthetic gene sequences from each line and assessed genomic structural variations (Supplementary Table 14). Multiple copies of the genes *CqDODA1* and *CqCYP76AD1*, which are involved in the conversion of L-tyrosine to betanine, are located at 74.53-74.71 Mbp, 74.60-74.82 Mbp, and 69.74-69.91 Mbp on chromosome 1B, and at 58.22-58.33 Mbp, 57.36-57.44 Mbp, and 52.04-52.10 Mbp on chromosome 2A in J075, J100, and QQ74, respectively. Additionally, chromosomes 4A and 4B had clusters of *CqDODA1*s (Figure 9A). In particular, regions on chromosomes 1B and 2A containing clusters of betalain synthesis genes were less conserved in synteny compared to adjacent regions in each lineage. Several genes, such as *CqCYP76AD1* on 1B, *CqDODA1* on 4A, and *CqDOPA5GT* on 4B, were more duplicated in the J075 and J100 lines than in the lowland QQ74 line. The leaf color at the shoot apex of the highland J075 and J100 lines was redder compared to that of J082, which is the inbred line derived from PI614886, the same accession as QQ74 (Mizuno et al., 2020; Ogata et al., 2021). Comparing the two highland lines, J075 displayed a darker red color at the shoot apex leaves and hypocotyl (Figure 9B). These color differences may be linked to variations in genomic sequences in regions harboring betalain biosynthesis genes.

**FIGURE 9.**
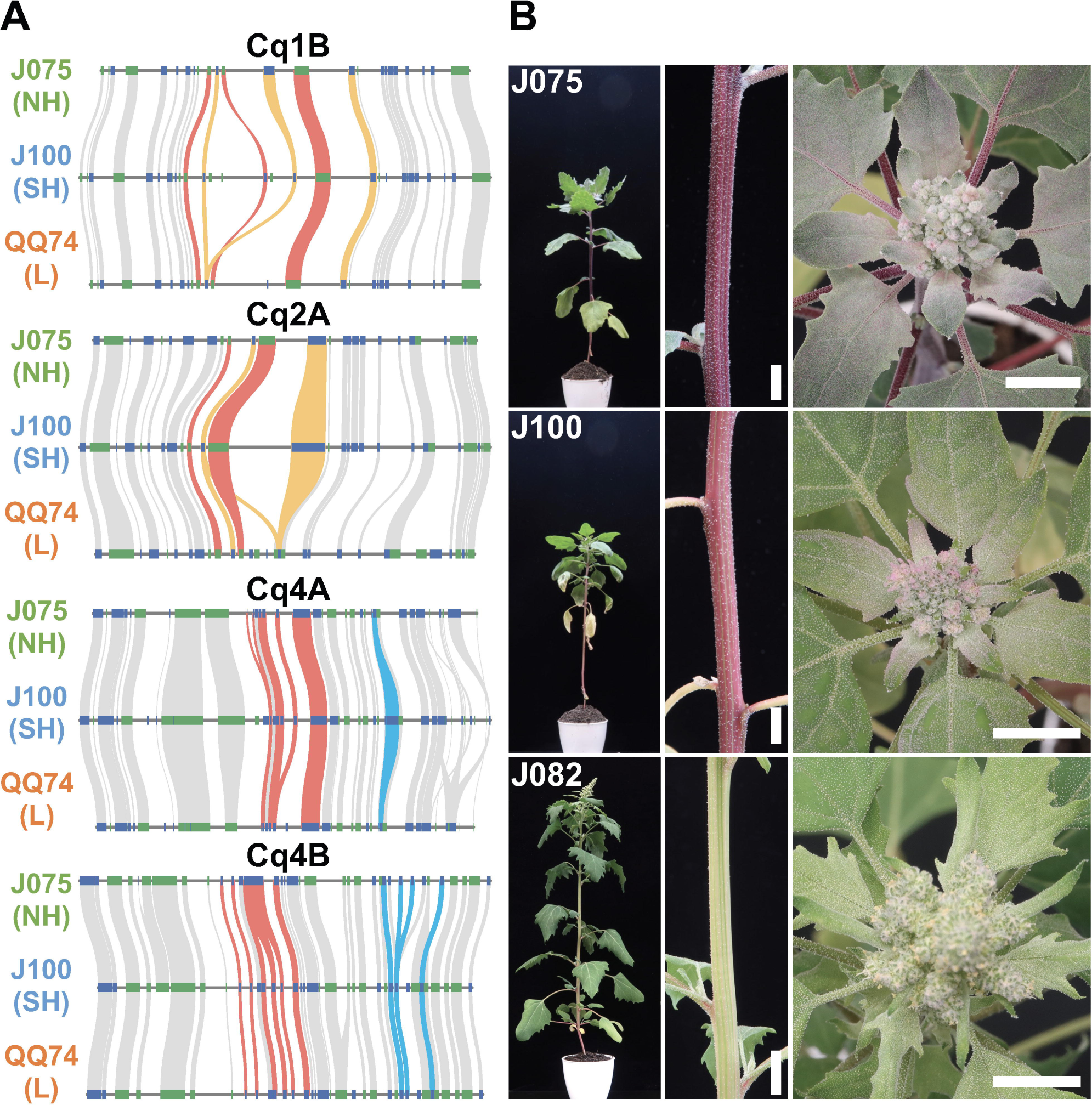
Syntenic betalain biosynthesis gene clusters and plant color phenotypes in representative quinoa lines. **(A)** Synteny analysis of betalain biosynthesis genes on chromosomes 1B, 2A, 4A, and 4B among J075, J100, and J082. The red, orange, and blue lines highlight the syntenic gene pairs *CqDODA1*, *CqCYP76AD1*, and *CqDOPA5GT*, respectively. NH, SH, and L indicate northern highland, southern highland and lowland, respectively. **(B)** Phenotypes of stem and shoot apex colors in J075, J100, and J082 after 56 days of growth. J082 is the inbred line derived from PI614886, the same accession as QQ74. Bars indicate 1 cm.

## 4 Discussion

Here, we present chromosome-scale high-quality *de novo* genome assemblies for two quinoa inbred lines from the northern and southern highlands of the Andean Altiplano of South America, where quinoa originated, using a combination of PacBio long-read sequencing with the dpMIG-seq method (Nishimura et al., 2024). Because quinoa can partially outcross due to the presence of both hermaphrodite and female flowers on the same plant (Maughan et al., 2004; Christensen et al., 2007), high-quality quinoa inbred lines need to be used for detailed molecular analysis, and high-quality genomes of highland quinoa lines grown close to their place of origin are essential for understanding the origin of quinoa and the process of domestication. These quinoa genomes of northern highland line J075 and southern highland line J100 are 1.29 Gb and 1.32 Gb, respectively, of which approximately 98.9% (1.28 Gb, Table 1) and 96.4% (1.27 Gb, Table 1) can be anchored to 18 chromosomes. The quality of these two genome assemblies of highland quinoa lines was higher than that of several other published quinoa genomes, including those of Kd (Yasui et al., 2016), QQ74 V1 (Jarvis et al., 2017), Real (Zou et al., 2017), CHEN125 (Bodrug-Schepers et al., 2021), and QQ74 V2 (Rey et al., 2023). It is worth noting that 72.6% and 71.5% of the repetitive element and 99.2% and 99.1% of the plant single-copy orthologs were detected in the assembly genomes of J075 and J100, which is a higher percentage than detected in Kd, QQ74 V1, Real, CHEN125, or QQ74 V2. Taken together, the assemblies of J075 and J100 are relatively accurate and complete. These are the most recent reference genomes for quinoa and will lay the foundation for understanding the expansion of quinoa cultivation areas and adaptation during intraspecies diversification in harsh environments and will provide important resources for the further investigation of genetic diversity in quinoa and related species.

The expansion and contraction of orthogroup gene families have contributed to plant diversification during evolution (Demuth and Hahn, 2009). Among these gene families, those related to nucleic acid metabolism and defense responses were particularly abundant in the genome of quinoa lines (Figure 5 and Supplementary Tables 10 and 11). Transposons, related to nucleic acid metabolism, have been identified as important in adaptive evolution (Li et al., 2018), suggesting that the expansion of these gene families was necessary for the adaptive evolution of quinoa populations, which are distributed in widely differing environments. Defense-related gene families have also been reported to expand through tandem duplication (Hanada et al., 2008), with the occurrence of multiple genes with similar functions possibly leading to diversification among quinoa lines. Beyond these common gene families, we identified features from gene families specific to each line, presumed to reflect the unique characteristics of the quinoa populations (Figure 5 and Supplementary Tables 10 and 11). The expansion of the auxin-related gene family in J100, and the post-embryonic development gene families in both J100 and QQ74, may be linked to enhanced post-germination growth in lines from the southern highland population compared to other populations (Mizuno et al., 2020). Furthermore, the expansion of gene families related to photosynthesis in both J075 and QQ74 suggests that variations in photosynthesis-related genes have evolved to adapt to the significant elevational and latitudinal differences in the regions where they are distributed.

Whole genome replication events, inferred from synonymous substitution rates at homoeologous genes between each line’s genome and progenitor genomes, confirmed that the same events occurred as those inferred from phylogenetic relationships within the species (Figure 6). Additionally, genomic structural variations were found between the southern and northern highland lines (Figure 8 and Supplementary Table 12). We have reported that agronomic traits and environmental stress responses are associated with quinoa genetic populations (Mizuno et al., 2020). In quinoa, plant color, derived from betalain, is one of the major phenotypic traits. We observed color differences among the highland lines J075 and J100, and the lowland line J082 (Figure 9B), which is the inbred line derived from PI614886, the same accession as QQ74 (Mizuno et al., 2020; Ogata et al., 2021). A comparison of gene-level synteny suggested that the highland lines J075 and J100 retained more copies of highland-type betalain biosynthesis genes in clustered regions of chromosomes than the lowland reference line (Figure 9). The color differences among lines should be considered in light of the low conservation of these regions and the influence of other genes in the betalain biosynthetic pathway. Gene families associated with structural variations may lead to the various phenotypic changes observed among quinoa populations. The comparative genomic analysis platform for lines J075 and J100, representing the northern and southern highland populations, has the potential to expand our understanding of genetic diversity among quinoa populations and will accelerate the identification of genes associated with phenotypes.

## Data availability statement

Raw data from this study were deposited in the DDBJ Sequence Read Archive database under the BioProject ID: PRJDB18071. The genome sequence data (PacBio and Illumina) are available under accession numbers DRR551518 - DRR551710.

## Author contributions

YK: Conceptualization, Data curation, Formal analysis, Funding acquisition, Investigation, Methodology, Visualization, Writing – original draft, Writing – review & editing. HH: Data curation, Formal analysis, Funding acquisition, Investigation, Methodology, Writing – review & editing. KS: Data curation, Formal analysis, Funding acquisition, Investigation, Methodology, Writing – review & editing. KN: Formal analysis, Funding acquisition, Investigation, Methodology, Writing – review & editing. KF: Investigation, Writing – review & editing. RO: Conceptualization, Project administration, Writing – review & editing. GA: Conceptualization, Project administration, Writing – review & editing. YN: Funding acquisition, Investigation, Project administration, Resources, Visualization, Writing – review & editing. YY: Conceptualization, Data curation, Formal analysis, Funding acquisition, Investigation, Methodology, Project administration, Writing – review & editing. YF: Conceptualization, Funding acquisition, Project administration, Resources, Supervision, Validation, Visualization, Writing – original draft, Writing – review & editing.

## Acknowledgments

We thank the staff of JIRCAS, M. Toyoshima, N. Hisatomi, Y. Masamura, K. Ozawa, Y. Shirai, N. Saito, Y. Nakamura, Y. Nonoue, Y. Takiguchi, A. Aoyama, Y. Ishino, N. Seko, T. Nada, and M. Nozawa for their excellent technical assistance, T. Ogata, Y. Murata, and T. Kashiwa for their useful discussion and support, the staff of the Kazusa DNA Research Institute, Y. Kishida, C. Minami, K. Ozawa, H. Tsuruoka, and A. Watanabe for skillful technical support, Y. Tokura, K. Katsura, I. Molares, A. Bonifacio, W. Rojas, M. Patricia, J. Quezada, Y. Flores, J. Tanaka, S. Kushino for their cooperation in quinoa research. Computations were performed in part on the NIG supercomputer at ROIS National Institute of Genetics.

