## Supplementary figures and images for "Chromosome-level genome assemblies for two quinoa inbred lines from northern and southern highlands of Altiplano where quinoa originated"

### Supplementary Figure 1

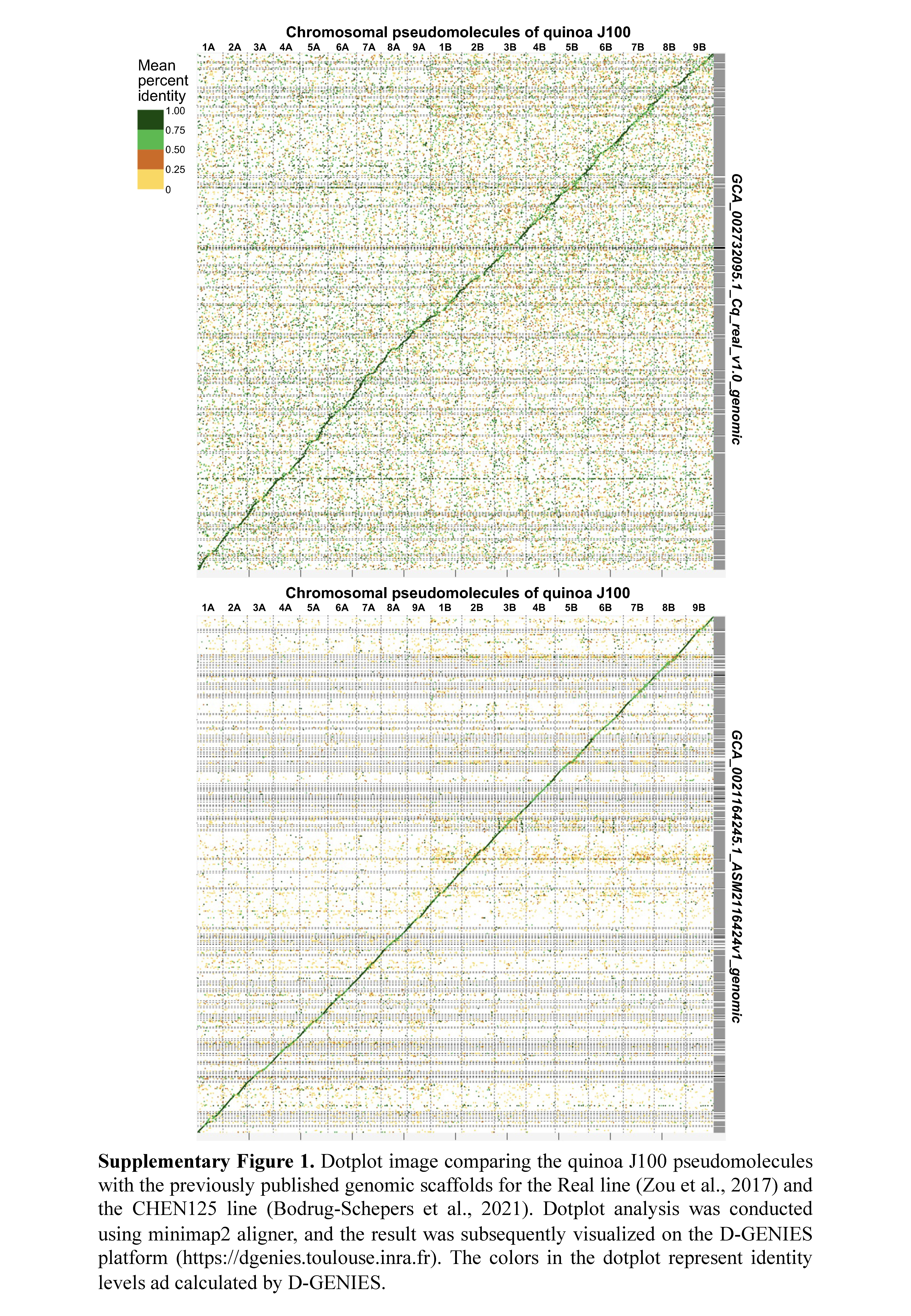

### Supplementary Figure 2

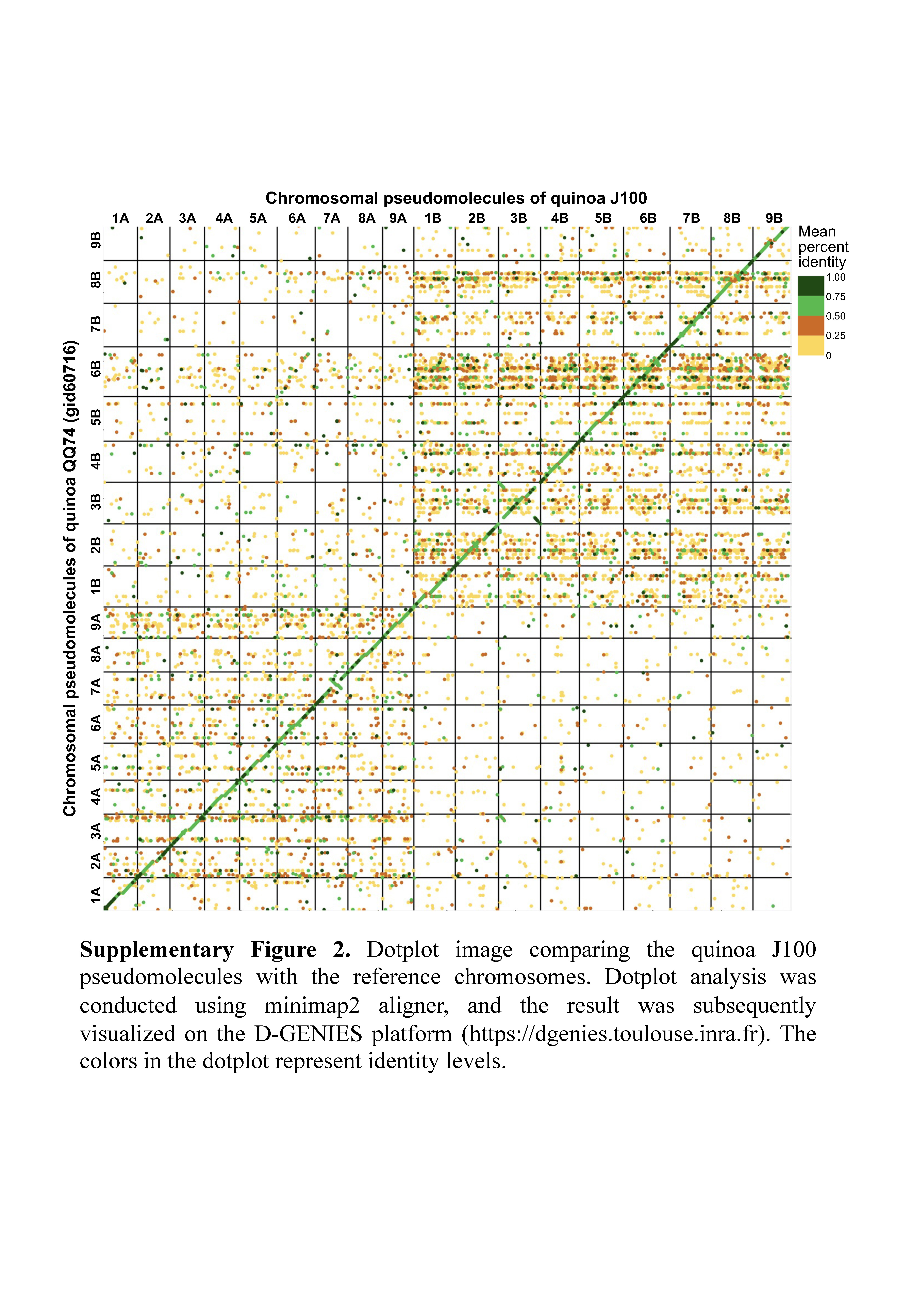

### Supplementary Figure 3

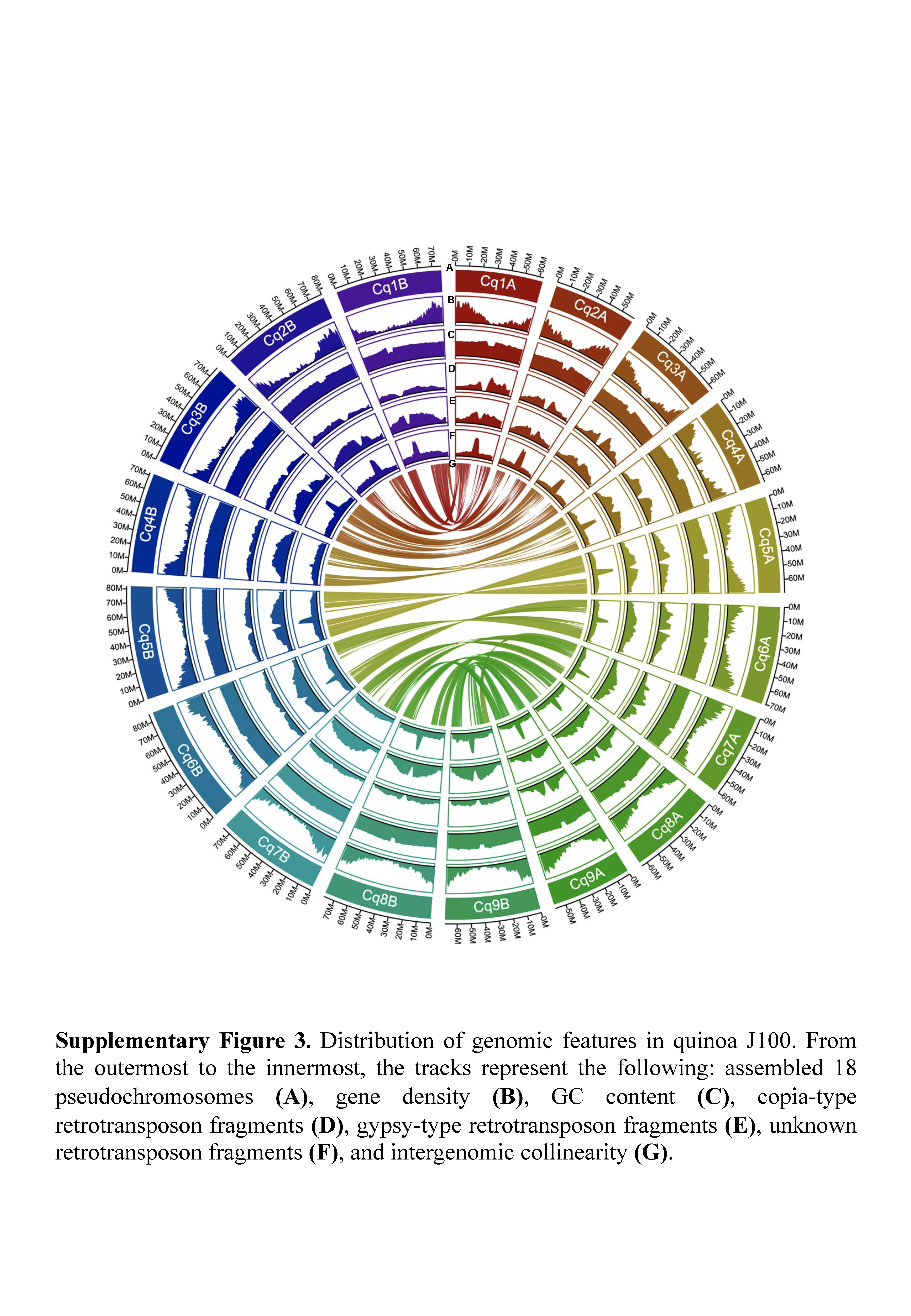

### Supplementary Figure 4

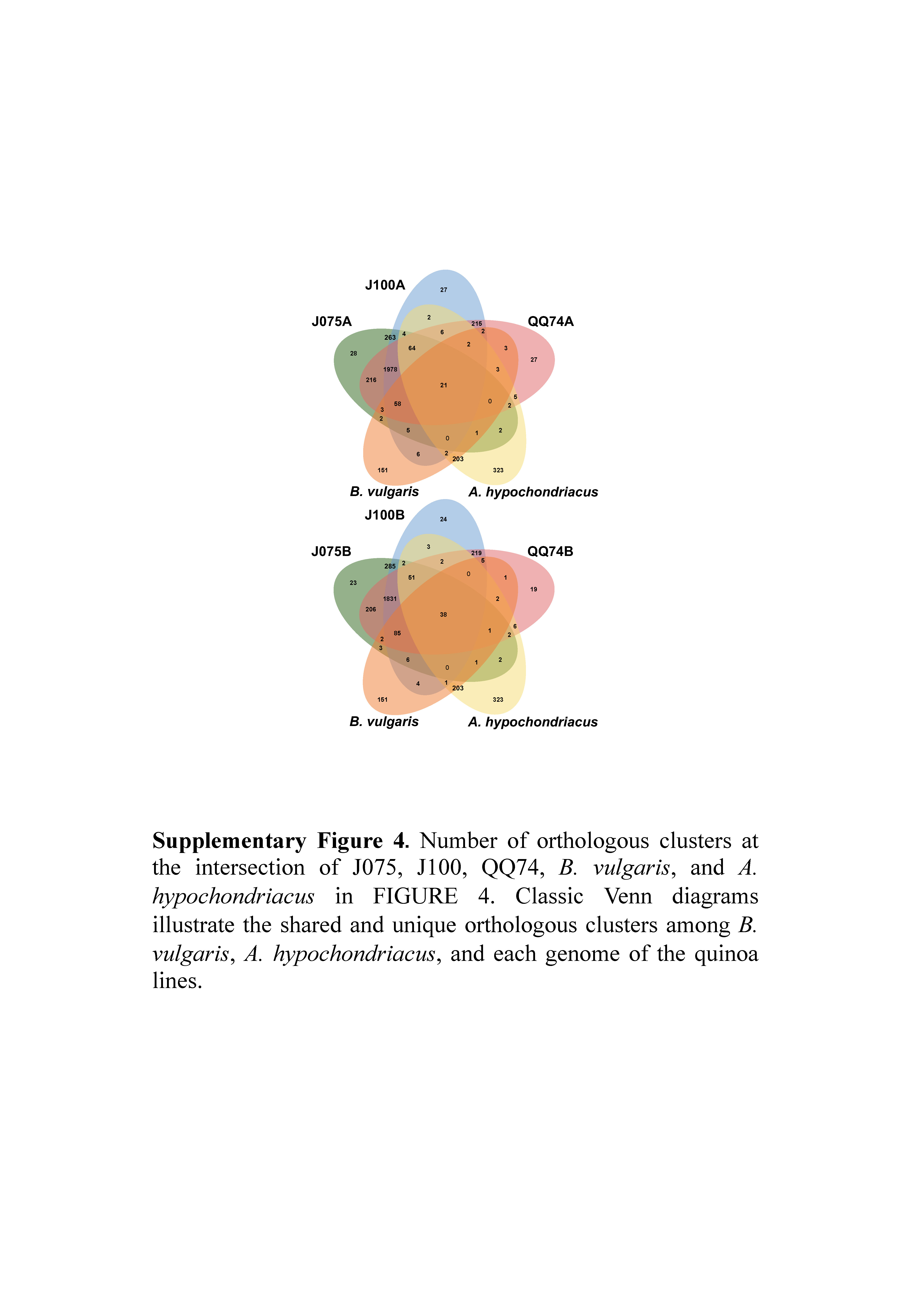
